# Local and regional scale mycorrhizal network assembly in an experimental prairie-pasture system

**DOI:** 10.1101/2022.10.05.510876

**Authors:** Katherine L. Shek, Hilary Rose Dawson, Toby M. Maxwell, Barbara Bomfim, Paul B. Reed, Scott Bridgham, Brendan Bohannan, Lucas C.R. Silva

## Abstract

Arbuscular mycorrhizal (AM) symbioses between plants and fungi are essential to the functioning of terrestrial ecosystems through maintaining soil stability, controlling nutrient cycles (e.g. C, N, P and K), and influencing competitive dynamics in plant communities. Despite the importance of AM symbioses, the ecological coassembly patterns of AM fungi-plant partners are not well characterized across environmental gradients. Further, it is unclear whether fungi forming associations with several plants of the same or different species – forming common mycorrhizal networks (CMNs) – preferentially allocate limiting resources within natural plant communities at the local-scale. We used an experimental prairie-pasture grassland system in three sites along a latitudinal gradient ranging from cool/wet to warm/dry climates to investigate how environmental conditions, local plant diversity and drought shift AM fungal composition and plant-fungal coassembly patterns across spatial scales. We show that plant-AM fungal assembly patterns are hierarchically structured, with environmental variables driving differences in AM fungal communities at the largest spatial scale (across sites), and plant host identity and diversity governing AM assembly at the local scale (within plot). Bipartite interaction networks revealed evidence for preferential partner selection between plants and fungi, while there was no evidence for nested assembly of plant-fungal partners. At the plot-level, we applied stable isotopes (^13^C and ^15^N) to illustrate CMN assembly and nutritional function. There was no significant correlation between increased resource transfer among plants in a plot that shared more AM fungal partners; however, we identified specific AM fungi that were indicator taxa for increased plant isotope enrichment. Further research integrating stable isotope probing of fungal DNA in plant roots is necessary to more clearly illustrate the form and function of CMNs in grasslands under different environmental and plant diversity conditions.

## INTRODUCTION

The arbuscular mycorrhizal (AM) mutualism is a symbiotic relationship between soil-dwelling fungi and >80% of vascular plants, and is key to the functioning of terrestrial ecosystems. In AM partnerships, communities of mycorrhizal fungi colonize the root systems of plants, and reciprocal exchange of nutrients and other resources occur in the rhizosphere^1^. The essential ecosystem services provided by these plant-fungal symbioses include increased resilience of communities to biotic and abiotic pressures, below-ground resource partitioning, and alteration of competitive dynamics in plant communities through equalizing fitness differences and expanding niche space. AM fungi are obligately bound to their plant hosts and serve nutritional functions through uptake and transfer of limiting nutrients such as N, P and K, in addition to providing increased resistance to stressors such as drought^1^. Plants vary in their dependency on AM fungi, and individual plants can form associations with numerous AM fungal taxa in their roots. Likewise, an individual AM fungus, or multiple conspecific AM fungal individuals, can form associations with multiple host plants, forming mycelial connections between neighboring root systems, called common mycorrhizal networks (CMNs)^2^. These networks can connect the rhizospheres of the same or different species of plants, thus influencing plant community dynamics, primary productivity, and the response of ecosystems to drought pressures^3^. Despite several studies investigating the impacts of CMNs on plant composition and behavior, the form and function of arbuscular CMNs in natural systems remain poorly characterized. For instance, differential allocation of resources among coexisting plants by the CMN has been shown in flax and sorghum plants^4^, but there is also evidence for lack of discrimination between high- and low-quality partners by the CMN^5^. One relatively unexplored factor that may drive differences in CMN behavior across these systems is the composition and diversity of AM fungi present in experimental settings, and whether different groups of AM fungal taxa vary in their contributions to CMN functions.

To identify AM fungi forming CMNs and characterize the emergent outcomes of CMN-mediated functions in plant communities, we must gain a better understanding of the factors that shape plant-fungal co-occurrences in nature. AM fungi are widely regarded as host generalists^6^, as there are currently only 384 estimated species of AM fungi known to form associations with a vast diversity of host plants^7,8^. However, recent evidence suggests a role of preferential partner selection in the AM symbiosis, with some studies suggesting trait-based correlations between AM partners^9^ and others supporting matching life histories^10^. Nevertheless, these patterns may depend on the identity of neighboring plants in the community^11^. Together these findings highlight the need for further investigation of how plant community diversity structures AM fungal composition and diversity, which has been shown to feedback to maintain plant diversity and ecosystem productivity^12^. Further, the species pool of AM fungal taxa available to colonize host plants in a given environment is largely driven by abiotic factors such as soil properties, climate, latitude and disturbance history^13^, adding environmental context and disturbance intensity in addition to host plant identity as determinants of AM fungal-plant associations across different systems^14^.

Ecological network theory is a common approach to decipher interaction patterns driving plant-AM fungal cooccurrences in nature^15^. This theory posits that mutualistic interaction networks almost always show patterns of nestedness, where specialist taxa interact with a subset of the partners that generalists interact with^16,17^. Several studies to date have shown AM fungal-plant interaction networks with significantly higher nestedness than expected randomly^18–20^. The opposite of nestedness in plant-fungal assembly patterns would show interaction networks with high modularity, in which interacting species form distinct groups and preferential partner selection plays a major role determining the composition of AM fungi colonizing individual plant species^9,21^. Network theory metrics such as these can help elucidate the relative roles of environment and host filtering on AM fungal communities. These network analyses are not to be confused with determining CMN structure and function; while network theory is generally applied at larger spatial scales to tease apart plant-fungal assembly patterns, CMNs occur at the local or plot-level spatial scale and require more refined analyses of fungal-mediated resource sharing between plants under varying environmental conditions.

Pastures are common agricultural systems governed by AM connections, where AM fungi play essential roles in plant nutrition and productivity under variable environments^22,23^. Pastures in the Pacific Northwest of the United States (PNW) are ideal systems to investigate the roles of AM fungal composition and plant resource acquisition and sharing via CMNs, because they likely rely on AM fungal functions to maintain plant diversity and productivity in the face of N-limitations. The PNW is a Mediterranean climate zone that was historically dominated by diverse grasslands and prairies, but decades of land management resulted in conversion to less diverse pastures that are dominated by introduced grasses^24^. The PNW has experienced increased extreme drought frequency and intensity which has caused measurable declines in PNW plant cover, especially for annual forbs and exotic annual grasses^25^. PNW prairie and pasture systems therefore provide the opportunity to simultaneously investigate the relative roles of host plant diversity, identity and drought pressures on plant-AM fungal coassembly patterns, and the form and function of CMNs.

Here, we first characterized how differences in environmental conditions, plant diversity and drought influence AM fungal composition and plant-fungal interaction patterns at broad spatial scales. We then utilized ^13^C and ^15^N stable isotopes at the plot-level to illustrate CMN assembly and nutritional function across plant diversity and drought treatments. Specifically, we address two major questions regarding plant-AM fungal assembly and CMN structure in PNW prairie/pasture ecosystems: (1) what biotic and abiotic factors drive shifts in the cooccurrences of AM fungi-plant partnerships in pasture/prairie plants; and (2) do CMNs form and facilitate nutrient transfer in high-vs. low-diversity plant communities under drought? We hypothesized that AM fungal communities exhibit hierarchical spatial structuring, where shifts in latitude, climate and soil properties structure which AM fungi are present across sites and at large spatial scales, and biotic factors such as plant host diversity and identity are major determinants of AM composition at the local scale^13^. We predicted plant-fungal interaction networks would exhibit nestedness, with relatively greater nestedness in low diversity plots, and greater levels of preferential partner selection in high diversity plots due to more host filtering for specific fungal partners across plant functional groups in high diversity plant communities. Finally, we hypothesized that CMNs are formed by widespread generalist AM fungal taxa that associate with nearly all plants in a plot, while more specialist AM fungal-plant interactions do not correlate with resources shared. To address our aims, we characterized AM fungal composition in the roots of plants in experimentally manipulated plots representing high and low plant diversity treatments and drought versus control treatments, all spanning three sites along a 520km latitudinal gradient in the PNW.

## METHODS

### Description of sites and experimental treatments

We conducted this study across three sites spanning a 520km gradient of three Mediterranean climate combinations along the Cascade Mountains into the Siskiyou Mountains: cool and wet (western Washington), warm and wet (central Oregon), warm and dry (southern Oregon). The experimental plots used in this study consisted of 20 plots per site under 2 climate treatments: control and −40% of precipitation, with 5 replicate plots for each combination of diversity and climate treatments (Fig. 1). The drought plots had rain-out shelters installed in February 2016 that covered 40% of the plot area with clear acrylic plastic (MultiCraft Plastics, Eugene, OR, USA), which absorbed ≤8% of light transmittance^30^.Within each site, plots are divided into two plant diversity treatments: a high-diversity prairie-grassland state and a low-diversity pasture state. The high-diversity plots were planted using an identical mix of 29 native grasses and forbs seeded in 2016 to 2018 (more details in references 26-29^26–29^), while the low-diversity pasture plots were not manipulated and are dominated by one to a few species of introduced perennial pasture grasses (i.e. *Schedonorous arundinaceus, Agrostis capillaris, Alopecurs pratensis*). Over the course of the experiment, non-native species have also self-seeded into these plots, but overall manipulated plots have maintained higher plant diversity than the unmanipulated (hereafter, low diversity) plots which were not intentionally planted.

**Figure 1.**
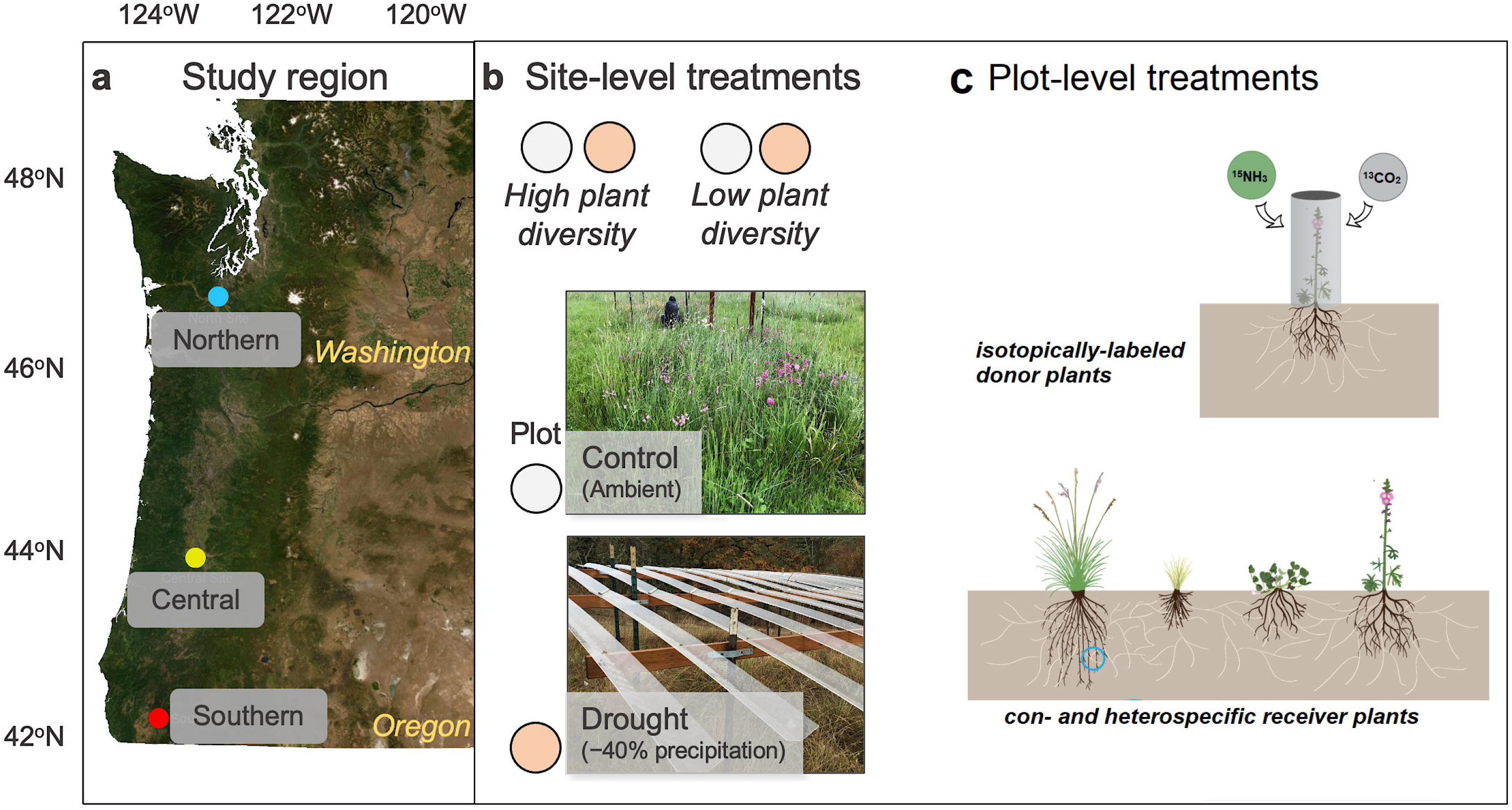
Experimental design for the study. Sites along 520km latitudinal gradient (a) from southern Oregon (warm and dry site, ‘Southern’), central-western Oregon (warm and wet, ‘Central’) and central-western Washington (cool and wet, ‘Northern’). Within each site, 5 plots of each treatment combination were sampled (high and low diversity, control and drought) for 20 plots per site (b). Within each plot, we sampled donor plants (labeled with ^13^C and ^15^N) and >6 other plants of the same and different species per plot (c). Figure adapted from Bomfim et al.^64^.

### Stable isotope labelling and plant harvest

Baseline samples were collected before labelling, between April 15 and May 20, 2019. To avoid destructive sampling of plants in the plots, we collected soil cores 0-10cm depth in triplicate for each plot. Root fragments were picked from soil core samples for DNA extraction and encapsulation for stable isotope analyses. We sampled baseline leaves for each plant functional group (annual/perennial, grass/forb) from biomass samples collected in 2018.

We selected plants located roughly in the center of each plot as ‘donor’ individuals that were *Sidalcea malviflora* in high diversity plots, and *Alopecurus, Schedonorus* or *Agrostis* in low diversity plots. The donor plants each received isotopically labeled ^13^CO_2_ and ^15^NH3 gas within a clear plastic cylinder at 20-minute intervals; labeled CO_2_ was pulsed three times enriched at 98 atm % ^13^C, and NH3 was pulsed twice at 98 atm % ^15^N. All labeling took place between 11am and 3pm on dates selected based on peak NDVI for each site, between April 22 and May 25, 2019. We selected ≥5 (average 10) additional plants of the four functional groups within each plot as putative ‘receivers’ at variable distances from donor plants, representing both con- and heterospecific plants.

We collected leaf samples were collected from all donor plants at the time of labelling, and of all donor and receiver plants at roughly 4- and 10-days post-labelling. At 21 days post-label, we harvested the entire plants and rhizosphere soils for all donor and receiver individuals into sterile WhirlPak bags and transported them on ice until storage at −20°C in the laboratory.

We processed all plants as soon as possible after collection. Plant processing consisted of shaking off rhizosphere soils into sterile WhirlPak bags for storage at −80°C, followed by tracing above- and belowground plant parts to ensure root samples were from target individuals. We discarded roots that could not be traced to aboveground biomass of each individual before separating above and belowground biomass samples. Approximately 10x ~3cm fine root samples were picked for DNA extraction, and another subset of roots were oven dried before encapsulating for stable isotope analysis at the UC Davis Stable Isotope Facility (Davis, CA, USA). Leaves were also oven dried before encapsulating for stable isotope analysis. The remaining belowground plant tissue was stored at −20°C until further analysis.

### AM fungal community composition

DNA was extracted from root samples representing 450 individual plants using Qiagen DNeasy Powersoil HTP kits (Qiagen, Hilden, Germany) following the manufacturer’s protocol. To characterize AM fungal composition in each sample, we adapted a novel two-step PCR protocol for preparation of Illumina libraries to amplify a ~550bp fragment of the SSU rRNA gene of AM fungal genomes, which is currently the most well-supported region of AM fungal rRNA for taxonomic resolution^31^ and amplicons can be queried against a well-curated database for taxonomic assignment ^7^. The primers we used for this region are WANDA (5’-CAGCCGCGGTAATTCCAGCT-3’) and AML2 (5’-GAACCCAAACACTTTGGTTTCC-3’)^32,33^. The first round of PCR amplified the target region using WANDA and AML2 primers modified to include heterogeneity spacers and Illumina TruSeq ‘stubs’ in 20uL reactions consisting of: 2uL genomic DNA template, 10uL GoTaq Green Master Mix (Promega Corp., Madison, WI, USA), 6.9uL nuclease-free water, 10uM of each primer and 0.1uL BSA. PCR1s were run on a BioRad T100 thermal cycler (BioRad, Hercules, CA, USA) for the following protocol: initial denaturation 94°C for 3min, followed by 30 cycles of 94°C for 45s, 50°C for 60s, 72°C for 90s, final extension 72°C for 10min. Amplification success and amplicon length were confirmed for all PCR1 products on 1% agarose gels before proceeding with PCR2. PCR2s were carried out in 25uL reactions containing 3uL undiluted PCR1 product as template, 1uL of each forward and reverse PCR2 primers, 10uL GoTaq Green Master Mix, 0.1uL BSA and 9.9uL nuclease free water. These reactions were run on the same thermal cyclers with the following conditions: initial denaturation at 94°C for 3min followed by 12 cycles of 94°C for 45s, 52°C for 60s, 72°C for 90s with a final extension at 72°C for 10min. PCR2 primers bind to the Illumina TruSeq ‘stubs’ on PCR1 products, and add Illumina adapters and indices for multiplexing several samples on a single sequencing lane. Our PCR2 primers have unique indices on both forward and reverse primers, permitting multiplexing several projects on a single run. Further details regarding primer modifications and PCR protocols provided in the Supplementary Information (Note S1).

Successful PCR2 amplicons were quantified using the Quant-iT PicoGreen dsDNA Assay Kit (Invitrogen, Waltham, MA, USA) on a SpectraMax M5E Microplate Reader (Molecular Devices, San Jose, CA, USA). Quantification data were used to pool amplicons in equimolar concentrations, before PCR purification with QIAquick PCR Purification kits (Qiagen). Purified pools were sequenced on the Illumina MiSeq platform (paired-end 300bp, Illumina Inc., San Diego, CA, USA) at the University of Oregon Genomics and Cell Chracterization Core Facility (Eugene, OR, USA). Sequence data are available through the NCBI Sequence Read Archive, BioProject accession number PRJNA####.

### Raw sequence processing and taxonomic assignment

All sequence data were demultiplexed and deduplicated using an in-house bioinformatics pipeline that handles the different aspects of our novel PCR protocol. Briefly, reads are re-oriented by querying each individual sequence for the biological primers before demultiplexing sequences into their respective samples with generous tolerance and quality thresholds, since downstream processing (see next steps, DADA2) performs sequence quality filtering. Orienting and demultiplexing scripts are in-house python scripts. Once sequence reads are assigned sample IDs, our pipeline utilizes the heterogeneity spacers in PCR1 primers as unique molecular identifiers (UMIs) that can parse apart duplicated sequences due to PCR bias as opposed to true biological multiplicity. The deduplication step aligns reads to the taxonomic reference database (MaarjAM7) using bowtie2^34^ and assigns UMI tags to each sequence using samtools^35^. UMItools is then used to filter out duplicate reads from PCR duplication (those with the same UMI), and retains reads with the highest alignment score to the reference database. Deduplicated sequences were used as input to the DADA2 pipeline for quality filtering and assembling into amplicon sequence variants (ASVs) with standard parameters^36^, which does not cluster sequences at a simialrity threshold like traditional OTU approaches, maintaining strain-level diversity in DNA sequences in our target region^37^. Taxonomy was assigned using the MaarjAM database with the ‘assignTaxonomy’ function in DADA^27^. Any ASVs that were not assigned taxonomy by DADA2 were searched using the MaarjAM BLAST+ feature in the gDAT bioinformatics pipeline^8^; we manually added these taxonomic assignments to our resulting ASV table if they met the following BLAST criteria: ≥95% alignment, a BLAST e-value <1e-50, and ≥90% sequence similarity. The MaarjAM database classifies AM fungal sequences into ‘virtual taxa’ (VT) as proxies for AM fungal species; here we perform analyses both at the ASV and VT levels.

To normalize differences in ASV counts across samples, we performed variance stabilization in the Deseq2 R package^38^. This method utilizes a Bayesian mixture model that scales ASV counts within and across samples, avoiding taxon abundance biases introduced by traditional rarefying methods^39^. We also retained the unnormalized dataset for some of the analyses (see below).

### Plant community composition and soil nutrient data

Plant community composition in the experimental plots were surveyed and studied extensively since initial seeding; for fine details for seasonal plant surveys see Reed et al.^27^. We used point-intercept methods were used to characterize plant composition at the plot-level, and these count data were used to calculate plant species diversity for each plot. We calculated observed richness (number of plant species in each plot), Faith’s phylogenetic diversity and Rao’s phylogenetic diversity using a phylogenetic tree pruned from a preexisting mega-tree using the ‘phylo.maker’ function in V.PhyloMaker^40^. Soil nutrient data were collected using buried anion and cation probes (ug 10cm^−2^ burial period ^−1^, Plant Root Simulator Probes, Western Ag, Saskatoon, SK, Canada) as described in Reed et al. ^27^.

### Statistical analyses

All statistical analyses were performed in R version 4.0.3, and all plots were generated using ggplot2 unless otherwise indicated^41^.

We analyzed AM fungal community diversity and composition using the *phyloseq* package^42^. To address our first hypothesis, we tested whether site, plant diversity treatment and drought treatment significantly correlated with shifts in AM fungal composition using PERMANOVA models and distance-based redundancy analysis (dbRDA) with vegan functions ‘adonis2’ and ‘capscale’, respectively^43^.

We calculated two different metrics for nestedness, as each metric represents the compositional nestedness of plant-AM fungal networks differently. First, we used the ‘nestedtemp’ function in the vegan package to calculate the temperature of presence-absence interaction matrices for high- and low-diversity plots separately^43^. Matrix temperature is a normalized value describing how far interaction data diverge from being perfectly nested; that is, how many interactions exist outside of the hypothetical presences and absences in a perfectly nested matrix^44^. A perfectly nested matrix would have a temperature of 0, while a nestedness temperature of 100 would be considered anti-nested. The other nestedness metric we measured was NODF (nestedness based on matrix overlap and decreasing fill) which is another quantitative measurement of nestedness that separately accounts for plants and fungi (rows and columns in interaction matrices, respectively) and standardizes differences in total interactions; NODF values are the opposite of temperature, where a NODF value of 100 is a perfectly nested matrix^45^. All nestedness calculations were repeated with a randomized version of the community matrix using ‘quasiswap’ with ‘oecosimu’ in *vegan* with 500 simulations. This function tests the null hypothesis that the nestedness statistic is significantly different from a series of simulated random communities while retaining row and column sums of community matrices^43^. To examine whether the anti-nestedness in plant-AM fungal interaction networks was due to greater specialization in partner interactions, overall network specialization (H2’, Bluthgen et al. 2006) was calculated.

We used the *bipartite* package^46^ to calculate network-level specialization (H2’)^47^ in addition to network modularity with the Beckett algorithm^48^. We performed all of these analyses on presence-absence data aggregated by plot type then plant species. We calculated modularity for plant species-AM fungal interaction networks separately for high and low diversity plots at both the ASV and VT level. Values for each network were calculated using the ‘metaComputeModules’ function with 100 simulations and tested for significance against null models with 100 simulations of randomized networks with the ‘r2dtable’ function, which keeps row and column sums constant but randomly changes cell values in an interaction matrix^49^. We compared observed modularity scores (Qobs) relative to the null distribution (Qnull) using z-scores:

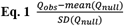

We performed threshold indicator taxa analysis (TITAN) to identify specific AM fungal ASVs that change in occurrence, frequency of occurrence, and direction of change with leaf ^15^N enrichment values for receiver plants only^50^. The ‘TITAN2’ function calculates indicator taxa values from traditional indicator species analyses with a continuous environmental variable and identifies change points along the environmental gradient at which individual ASVs change in abundance or frequency of occurrence. We bootstrapped these calculations and standardized them to z-scores, maintaining magnitude and direction of the ASV responses to environmental variables (in this case, leaf ^15^N enrichment). We transformed fungal ASV table to presence-absence before TITAN analysis. We visualized indicators using the ‘plotTaxaRidges’ function in TITAN2 ^50^.

To further examine the specific factors driving shifts in AM fungal composition across sites and plot types, we used a constrained ordination method (distance-based redundancy analysis, or dbRDA), where specific soil properties and plot-level plant species diversity variables were modeled against the Bray-Curtis dissimilarity matrix for AM fungal composition; the model included plant identity, soil nutrient data and plant species richness.

## RESULTS

After removing low-coverage samples (samples with fewer than 100 reads) we obtained 1,352 AM fungal ASVs belonging to seven families, eight genera and 95 virtual taxa in 429 samples. Of the ASVs detected in our dataset, 1,026 were in high diversity plots and 745 in low diversity plots. There were 10 ASVs that did not have taxonomy assigned in DADA2 and did not meet our BLAST+ criteria (<90% sequence similarity with taxonomy in MaarjAM database). Although our samples were not equally distributed across all host plant species, we observed saturation of accumulation curves for ASV richness for all plant species in both high and low diversity plots (Fig. S1).

### AM fungal diversity varies with latitude and plant diversity, but not drought treatment

We first examined whether AM fungal richness varied with plant functional group, lifespan, species, and plot-level diversity metrics. For plant species and functional groups, we detected significantly higher AM fungal ASV richness in perennial grasses compared to annual grasses in both high and low diversity plots (Fig. S2B, Wilcoxon p ≤ 0.015). Overall, there were no differences in AM fungal richness across plant species, but a pairwise post-hoc analysis revealed significantly lower fungal richness in *Vulpia* spp. and *Bromus hordaceus* compared to *Sidalcea malviflora* and *Schedonorous* spp. (Fig. S2A, Tukey’s Honest Significant difference corrected p ≤ 0.01). We did not detect any differences in AM fungal richness in drought versus control plots.

When we grouped ASV data at the plot level, high diversity plots had significantly greater observed fungal richness compared to low diversity plots (Wilcoxon p ≤ 0.0001), a relationship which correlated with plant community phylogenetic diversity across plots (Fig. 2A-C, R^2^=0.18, p < 0.001).

**Figure 2.**
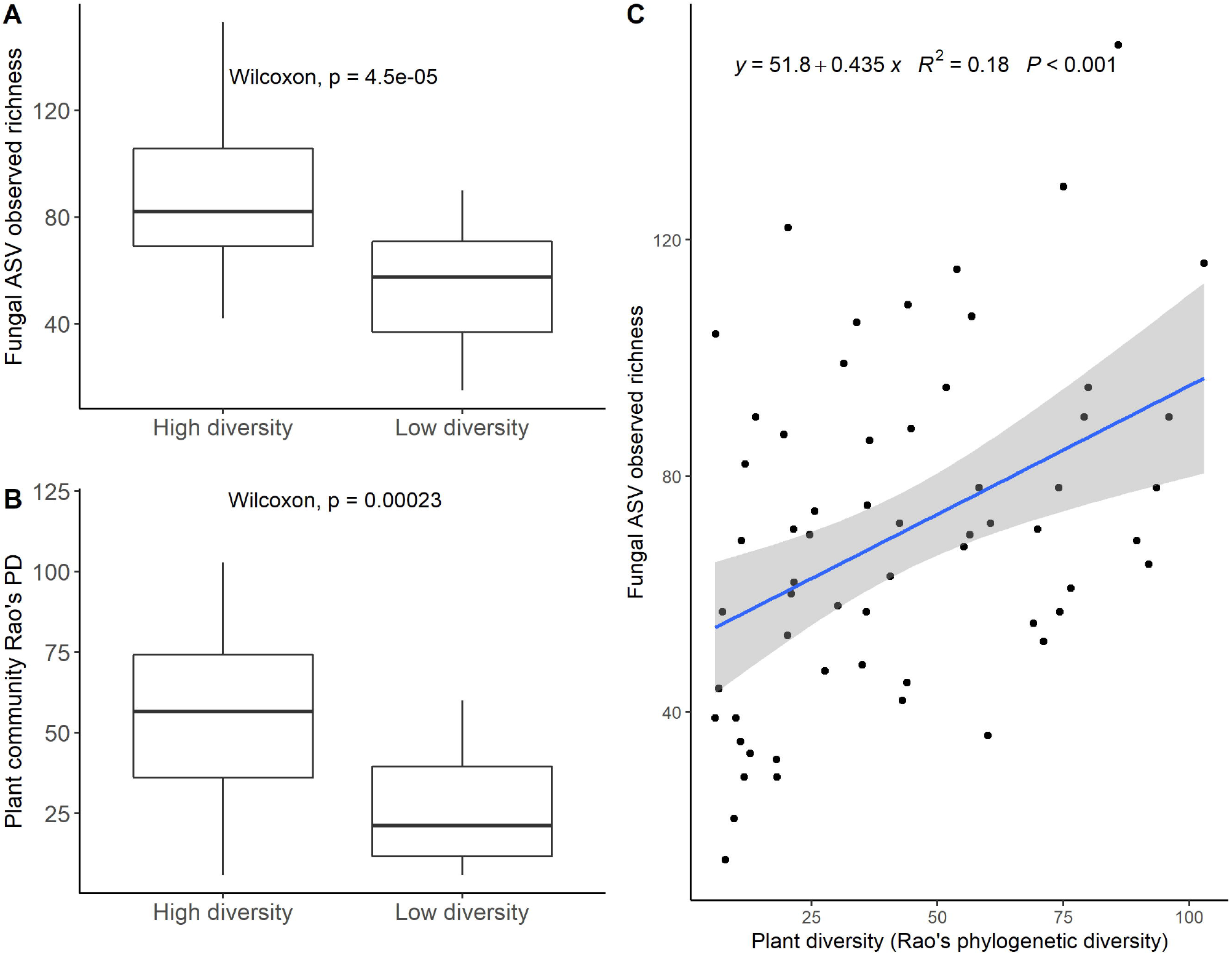
AM fungal alpha diversity (observed richness ASVs) reflected aboveground phylogenetic diversity patterns at the plot level. High diversity treatment plots had greater ASV richness (A) and phylogenetic diversity in the plant communities (B). The correlation between ASV richness and plant phylogenetic diversity was significant (C).

Sites had significantly distinct shifts in AM fungal composition as measured by Bray-Curtis dissimilarities at both the ASV and VT levels of AM fungal classification (Table 1, PERMANOVA p ≤ 0.001). This pattern remained significant when considering all plots, within each plot type, with an interaction between site and plot type (high vs low diversity treatments; Table 1, PERMANOVA p ≤ 0.001). Drought treatment did not affect AM fungal composition, except for weakly significant correlations at the VT level for the whole dataset and within pasture plots (Table 1, PERMANOVA p ≤ 0.01). Host plant species, plant functional group (grass vs. forb) and plant perenniality (perennial vs. annual) were all significant factors driving shifts in AM fungal composition, except for in the high diversity plots, where functional group was only moderately (ASV level) or not significant (VT level, Table 1).

**Table 1.**
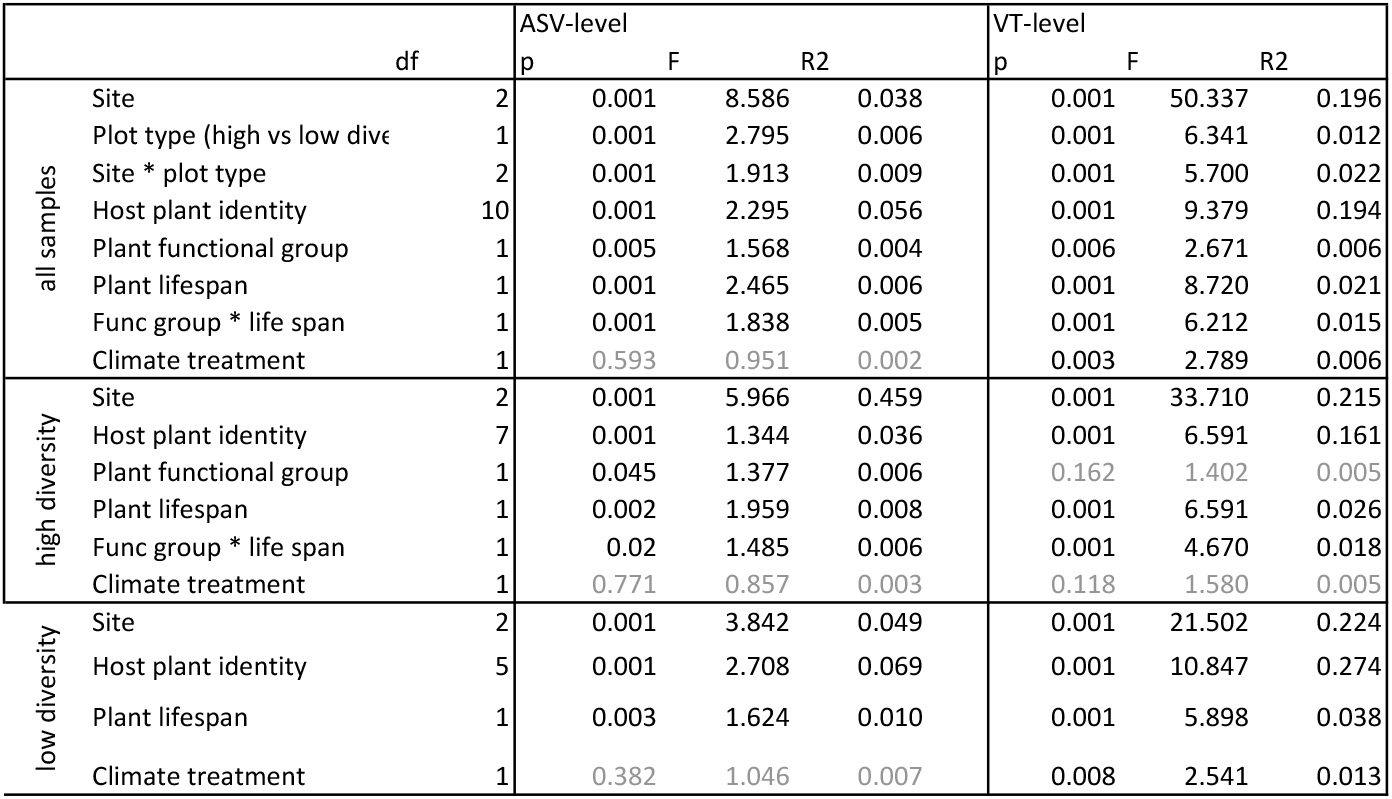
Results of permutational analysis of variance (PERMANOVA) models run on the Bray-Curtis dissimilarity matrices for normalized AM fungal sequence data at the level of ASV and VT. All models were run for each variable independently with 999 permutations except where an interaction is indicated. Non-significant relationships in grey.

The overall dbRDA model was significant (adjusted R^2^ = 0.14, p ≤ 0.0001) and revealed that several soil nutrient properties (total N, Ca, Mg, K, B, Fe, Mn, Cu, S, Al and Cd) as well as plant species richness were significant predictors of shifts in AM fungal composition (Fig. 3).

**Figure 3.**
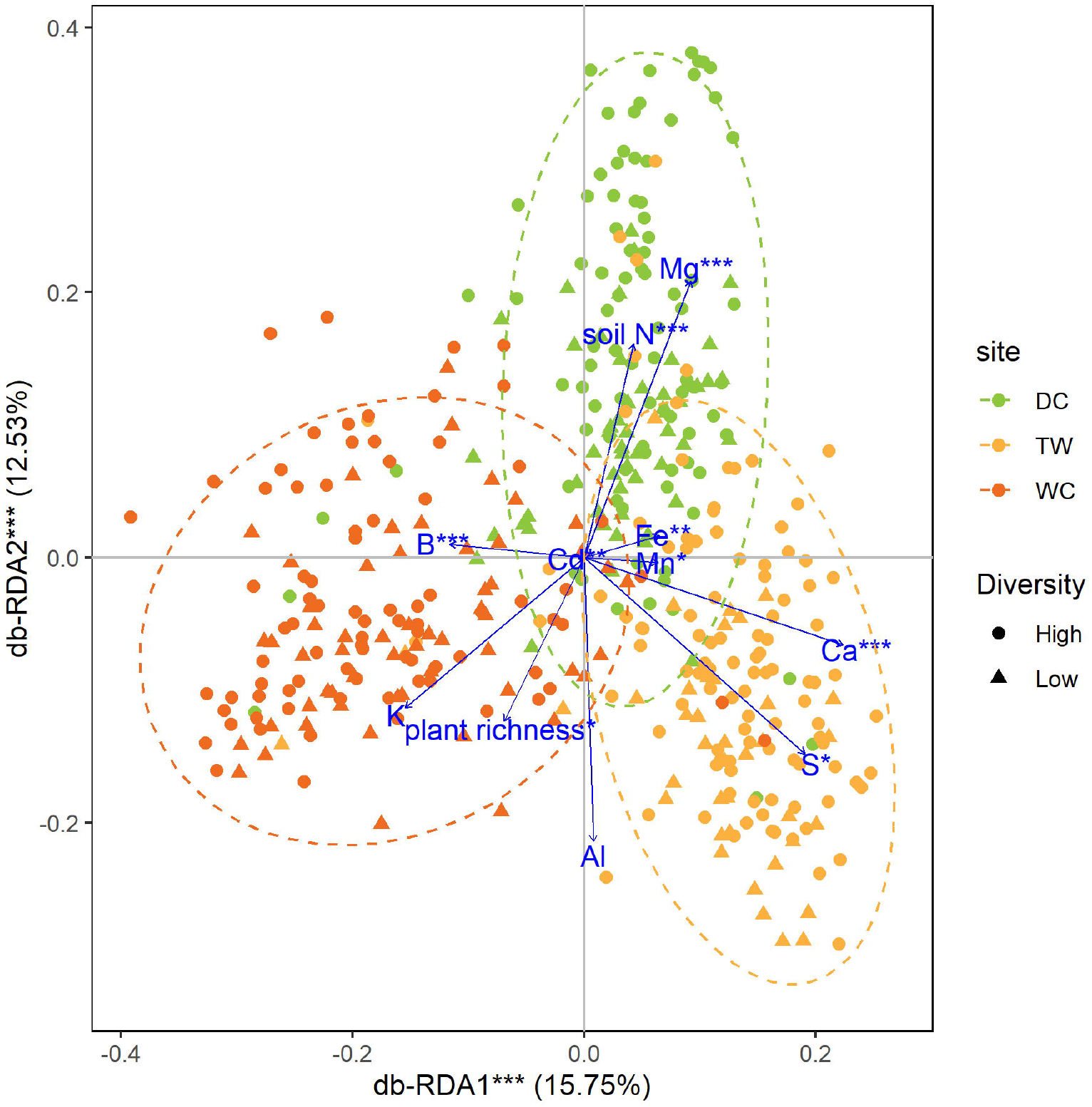
Constrained ordination analysis of Bray-Curtis dissimilarities for AM fungal communities at the ASV level across sites (colors) and plant diversity treatments (shapes). dbRDA model is significant (F = 1.5793, p ≤ 0.0001). Plot shows the main soil nutrient and plant richness variables correlating with shifts in AM composition (with 9999 permutations, * = 0.01 ≤ P < 0.05; ** = 0.001 ≤ P < 0.01; *** = P < 0.001; no asterisk = 0.05 ≤ P < 0.1). Dotted ellipses delineate 95% confidence intervals for grouping samples by site. DC = southern site, TW = northern site, WC = middle site.

### AM fungal-plant interaction network topology

Nestedness temperature for high diversity plots was significant relative to null models under 500 simulations (Fig. 4A; T=29.556, p ≤ 0.002), but not significant when using the NODF metric (NODF = 21.441, p=0.07). Low diversity plots had similar nestedness values, but did not significantly differ from null models for both temperature (Fig. 4B; T= 37.248, p=0.7126) and NODF (27.379, p=0.066) metrics. Overall network specialization was high. Low diversity plots had slightly higher metrics, but there was no difference between high and low diversity plot network specialization (p = 0.056, Fig. S4).

**Figure 4.**
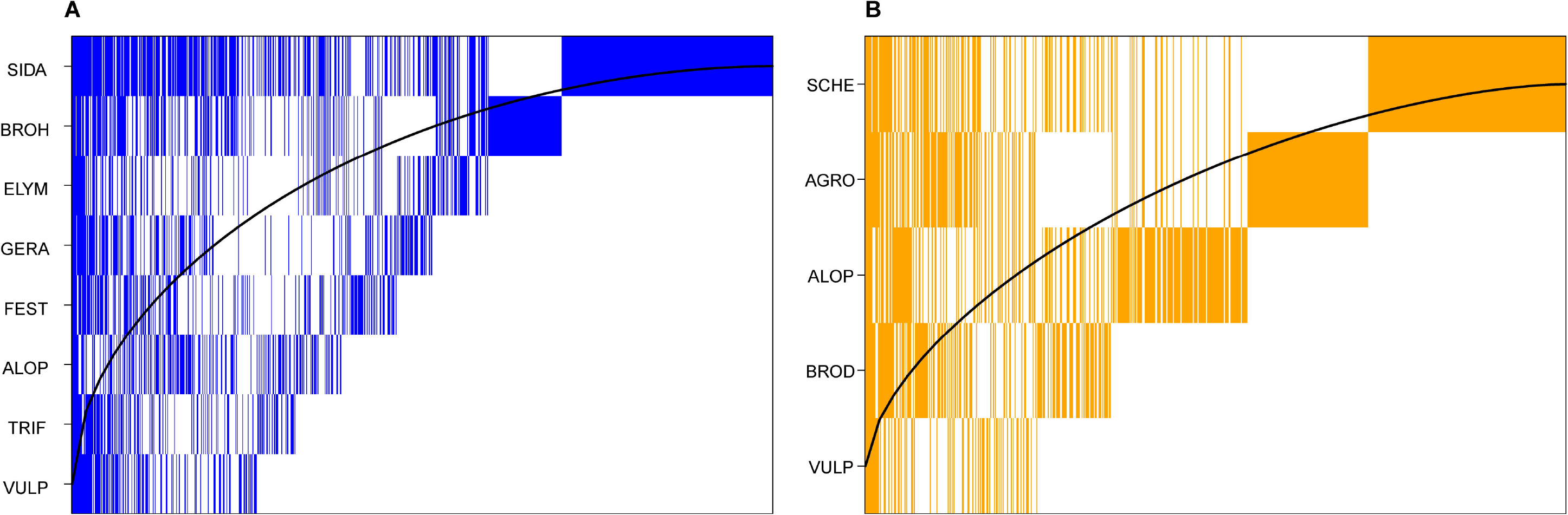
Nestedness of AM fungal ASVs (vertical lines) and their associations with host plant species (rows) for high diversity (A) and low diversity (B) plots across sites. High diversity plots had lower nestedness temperature (T=29.556) relative to the null models (p ≤ 0.002) but not when considering the nestedness based on overlap and decreasing fill of interaction matrices (NODF = 21.441, p=0.07). Low diversity plots were not significantly nested relative to null models. Curved lines show theoretical isocline where all interactions in a perfectly nested matrix would appear above the line (T=100 or NODF=0).

The AM-fungal-plant interaction networks were significantly modular relative to null models in both diversity treatments, both at the ASV and VT levels (Table 2, Fig. 5). All z-scores revealed observed modularity in networks ≥9 standard deviations more modular than random networks generated with the same marginal totals of interacting species (Table 2). ASV-level networks had significantly higher modularity values than the VT networks; due to the large number of ASVs in the ASV-level networks (>700), we present VT-level networks for visualization and more detailed analyses (Fig. 5). Low diversity networks were more modular than high diversity networks (Q_low_=0.2519, Q_high_=0.1453 for VT-level). Low diversity networks had three distinct groups of species interactions, whereas high diversity networks had four modules. Each distinct module consists of the plant-fungal interactions which occur significantly more frequently within the modules they belong to than with the taxa outside of the modules (Fig. 5). For each individual taxon’s membership in a module, among-module connectivity (c values) and within-module degree (z values) can be examined to better understand each taxon’s contribution to network topology^49,51,52^. Taxa with high values of both c and z are both major within- and between-module connectors and thus be considered major connectors in an interaction network^51^. We calculated c and z values for all AM fungi in both networks, to compare with subsequent local CMN and isotope enrichment analyses.

**Figure 5.**
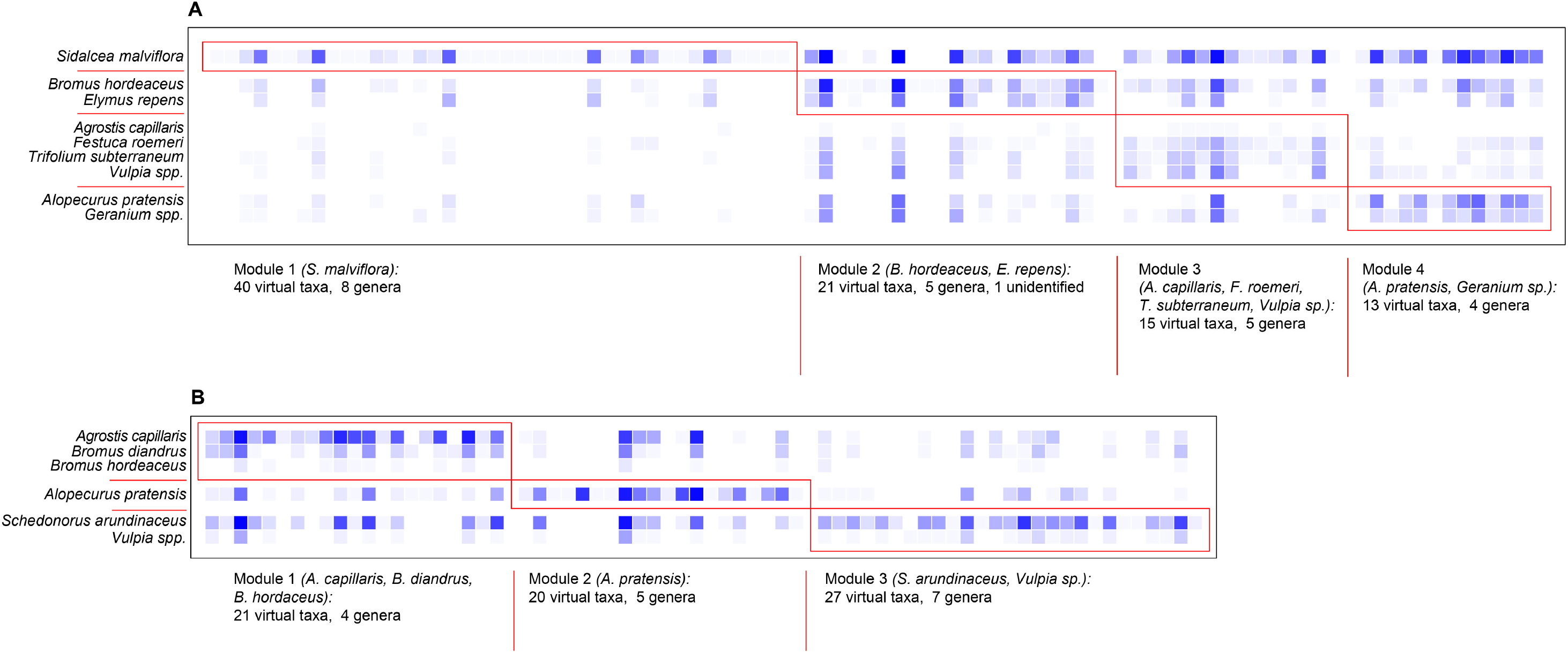
Plant-AM fungal interaction matrices with AM fungal VTs in columns and plant species in rows for high diversity (A) and low diversity (B) plots, ordered to show modularity and module membership of each taxon. Both matrices are significantly modular (high diversity Q=0.1453, p ≤ 0.0001; low diversity Q=0.2519, p ≤ 0.0001). Number of interactions for each plant-AM fungal VT combination indicated by color of tiles, ranging from 0 interactions (white) to 77 in high diversity and 39 in low diversity (brightest blue). Fungal VTs belonging to each module are listed in Tables S1 and S2.

### Plot-level network analysis and stable isotope enrichment

Leaf ^15^N enrichment was not related to the number of fungal ASVs shared between plants within a plot (linear mixed effects models p > 0.1). High diversity plots shared significantly more fungal ASVs between at least two plants (i.e., symbiont range of individual plants within a plot) compared to low diversity plots (Wilcoxon p ≤ 0.0001). We found significantly negative correlations between the first dbRDA axis and the number of shared fungal ASVs between plants in a plot, where the first axis largely discriminates between sites (Fig. S5, Fig. 3).

Threshold indicator taxa analysis (TITAN) detected 7 ASVs as significant indicators of increasing ^15^N enrichment in leaf samples (Fig. 6). This analysis calculates the classic IndVal index for indicator species analysis along a continuous environmental variable over several permutations and bootstraps, returning a standardized z-score representing the magnitude of IndVal scores for each fungal ASV that significantly increased (Fig. 6) or decreased (Fig. S6) with change in the leaf ^15^N enrichment variable.

**Figure 6.**
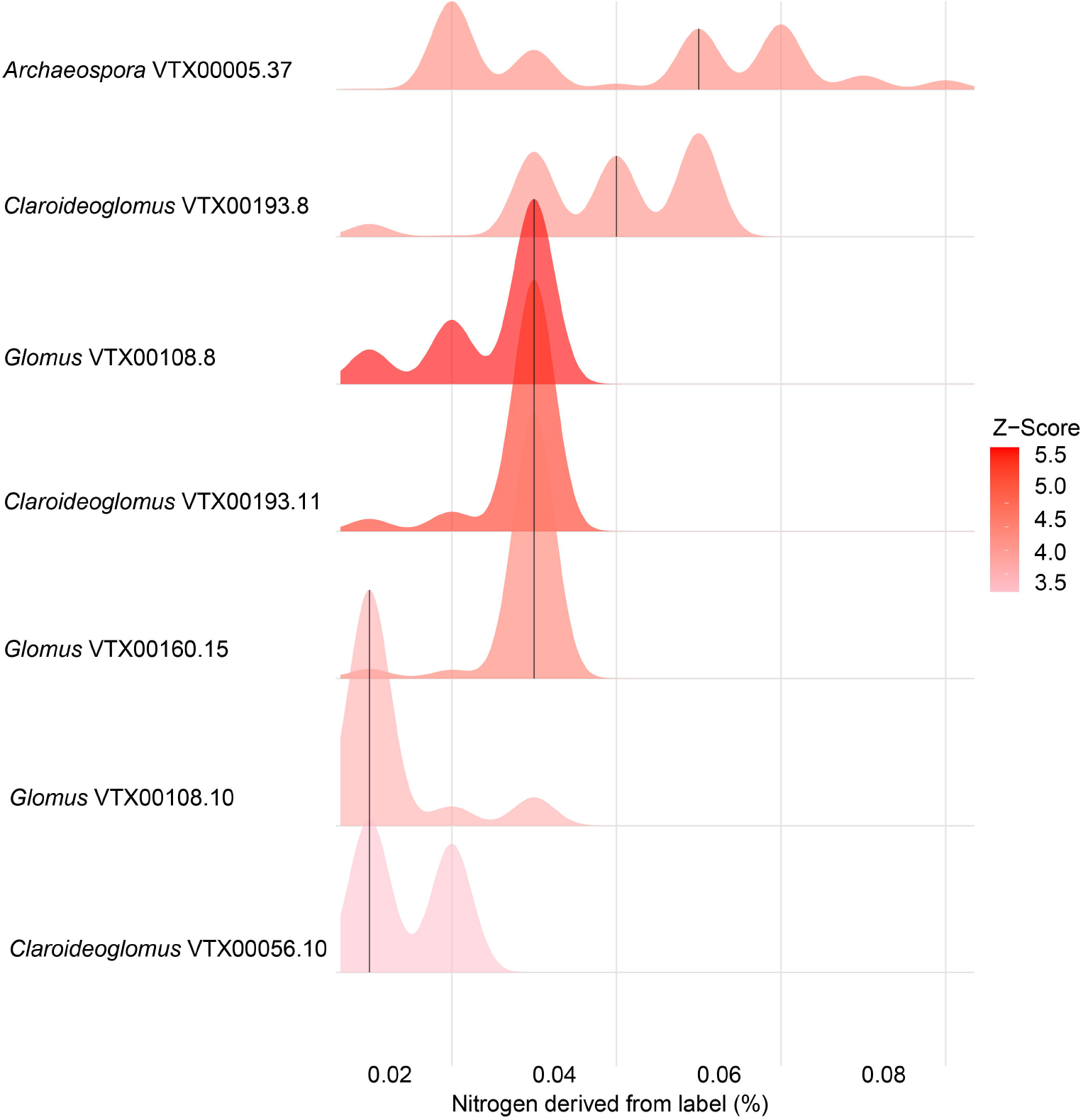
Indicator taxa analysis (TITAN) showing taxonomy for the AM fungal ASVs that increased in frequency of occurrences with increasing leaf 15N enrichment in host plants. Height of peaks indicate the magnitude of z-scores representing standardized IndVal scores (Dufrene and Legendre, 1997) bootstrapped 500 times with 250 random permutations.

## DISCUSSION

Theory suggests that AM fungi play essential roles in plant species coexistence, particularly in grasslands^53^. One major way in which AM fungi are thought to maintain plant coexistence in grassland ecosystems is through transfer of limiting resources, such as N, via CMNs^54^, but evidence for these behaviors in nature remains limited and we lack an understanding of how CMNs vary in structure and function across environments. We found that plant-AM fungal interaction networks in high and low diversity pasture systems display significant patterns of modularity and non-nestedness, implying greater levels of preferential partner selection than is expected in mutualistic networks. We also provided evidence for hierarchical spatial structuring in plant-AM fungal assembly patterns: across sites, AM fungal composition within roots of the same communities of host species was primarily driven by environmental variables at larger spatial scales; while within sites, plant host identity and local plant diversity shapes AM fungal assemblages at a more local scale. The absence of a significant relationship between number of shared AM fungal partners and amount of resources shared between plants at the plot-level suggests that there are more nuanced patterns of CMN-mediated resource transfer than frequency of interactions alone.

### Mycorrhizal fungal diversity increases with plant community diversity

We observed positive correlations between AM fungal richness and plant community diversity, both at the coarse level of plant diversity plot treatments, and within-plot phylogenetic diversity in plant communities. This finding is consistent with previous research and theoretical patterns in mycorrhizal diversity, in which aboveground diversity facilitates greater diversity in plant-associated microbial communities (reviewed in ref. 55 ^55^), including the abundance and diversity of mycorrhizal fungi ^56^. Specifically, different functional groups of plants have been shown to associate with distinct mycorrhizal fungal communities^57,58^, facilitating a consistent, positive relationship between plant and AM fungal alpha and beta diversity across biomes^59^. We found no evidence of an effect of long-term drought pressure on AM fungal community diversity across plant diversity treatments and sites. This finding was surprising, since AM fungi have been shown to respond strongly to drought in similar experimental grassland systems^60^, and generally provide plant community resilience under drought conditions^61,62^. This discrepancy in our study may be due to changes in plant-AM fungal behaviors, but not composition, under drought (i.e. plasticity of responses by both partners^63^), or differences in the severity of drought studied in other systems relative to ours. Due to the Mediterranean nature of our study system (temperate wet winters and hot dry summers), rainout shelters primarily affect plots only part of the year. This can reduce the observed drought effect to brief periods of intense rain where removing even 40% of received moisture is not enough to alter the conditions experienced by plants and fungi. Our finding of no effect is consistent with other studies published from this experiment (see Reed et al. 2019, Dawson et al., *in prep*)

### Mycorrhizal network assembly is shaped by environment and plant diversity

It is well-established that AM fungal composition is sensitive to differences in climate, soil properties and nutrient availability^64^, but few studies have addressed these patterns in nature at the community-level including multiple trophic levels. Our study analyzed how these factors shape both plant and fungal cooccurrences in both broad-scale interaction networks and local-scale common mycorrhizal networks. Unexpectedly, we found no evidence of nestedness in plant-fungal interaction networks, which is contrary to what is expected in mutualistic networks and shown previous studies if plant-AM fungal systems^16,18,20^. This result may be due to the finer taxonomic resolution at which we analyzed nestedness in interaction networks: the fungal ASVs in our study are considered distinct individuals with a single nucleotide difference in the small subunit (SSU) marker region of AM fungal genomes, while previous studies reporting significant nestedness in plant-AM fungal networks have examined patterns at coarser taxonomic scales (i.e. clustered sequences or genera). Nonetheless, the high modularity in our interaction networks suggest non-random interaction patterns between AM fungi and host plants in nature, a finding which is supported by previous studies examining host specificity in AM fungal communities^9,59^. Most of this preexisting research has been in relatively simple experimental systems either in the greenhouse or with single or few species consortia of mycorrhizal fungal taxa; our study provides one of the first characterizations of plant-AM fungal assembly patterns with naturally-occurring AM fungal communities under field experimental conditions of plant diversity manipulation. Our findings provide promising evidence that preferential partner selection in the AM symbiosis extends beyond simple experimental systems, and that even with the vast diversity of AM fungal taxa in natural soils (here, >1000 sequence variants and >90 virtual taxa), some level of specificity is apparent.

### Nutrient sharing via local common mycorrhizal networks

A major gap in our knowledge regarding CMN form and function is regarding whether the number of shared AM fungal taxa on roots of neighboring plants correlates with amount of resources shared through common AM fungal symbionts. We tested this using stable isotopic labeling and fine-scale AM fungal symbiont characterization (strain-level variation/ASVs) and found no evidence for this quantity of partners/quantity of resources relationship. To our knowledge, this has not been previously tested in complex, multi-species natural communities using stable isotope methods. To investigate whether resource sharing in CMNs is via higher quality partners rather than quantity of partners, we performed threshold indicator taxa analysis to get a general look at whether any AM fungal ASVs on plant roots served as indicators of stable isotope enrichment. The indicator ASVs selected by this algorithm were *Glomus*, *Claroideoglomus* and *Archaeospora* species, and the virtual taxa were all detected as ‘connectors’ (high c values) in our modularity analysis. This is theoretically interesting, as ‘connectors’ of modules in larger scale interaction networks implies that those taxa are linking distinct sets of interacting species, suggesting that resource sharing indicator taxa are connecting distinct groups of plants. Ecologically, this result may imply that loss of these species would cause a decline in CMN functioning in prairie/pasture systems in the PNW. Further research is necessary implementing stable isotope probing in fungal DNA extracts, to confirm whether these indicator AM fungal taxa are actively transporting isotope-enriched nutrients, or if they are simply indirectly responding to another factor that is driving CMN resource transfer functions.

## Acknowledgements

We would like to thank the Siskiyou Field Institute, The Nature Conservancy, and Capitol Land Trust for providing the sites for this experiment. Additional thanks to Laurel Pfeifer-Meister, Bitty Roy, Bart Johnson, Graham Bailes, Aaron Nelson, and Matthew Krna for their contributions to experimental design, and acknowledgements to several others for assistance with the HOPS project. This experiment was funded by National Science Foundation Macrosystems Biology grant #1340847, Plant Biotic Interactions grant #1758947, and Convergence Accelerator Pilot grant #1939511.

## Author Contributions

All authors designed the research; KLS, HRD, TMM, and PBR collected data; KLS and LCRS analyzed data; all authors wrote the paper.

## Notes

### Competing Interest Statement

The authors have declared no competing interest.

