## Supplemental information for "Local and regional scale mycorrhizal network assembly in an experimental prairie-pasture system"

### SUPPLEMENTARY INFORMATION

#### Note S1. Description of two-step PCR protocol.

Our two-step PCR protocol contains primer modifications and bioinformatics processing steps that account for some of the most pernicious issues that compromise modern high-throughput amplicon sequencing datasets, particularly those with longer amplicon lengths. The major factors of Illumina sequencing that our protocol accounts for include: poor reverse sequence read quality, which affects the merging of paired-end reads during bioinformatics processing of sequences; low base-call diversity in amplicon-generated datasets due to conserved regions of marker genes; PCR amplification biases that often result in inaccurate estimates of sequence abundances and primer interactions with Illumina adapters and indices.

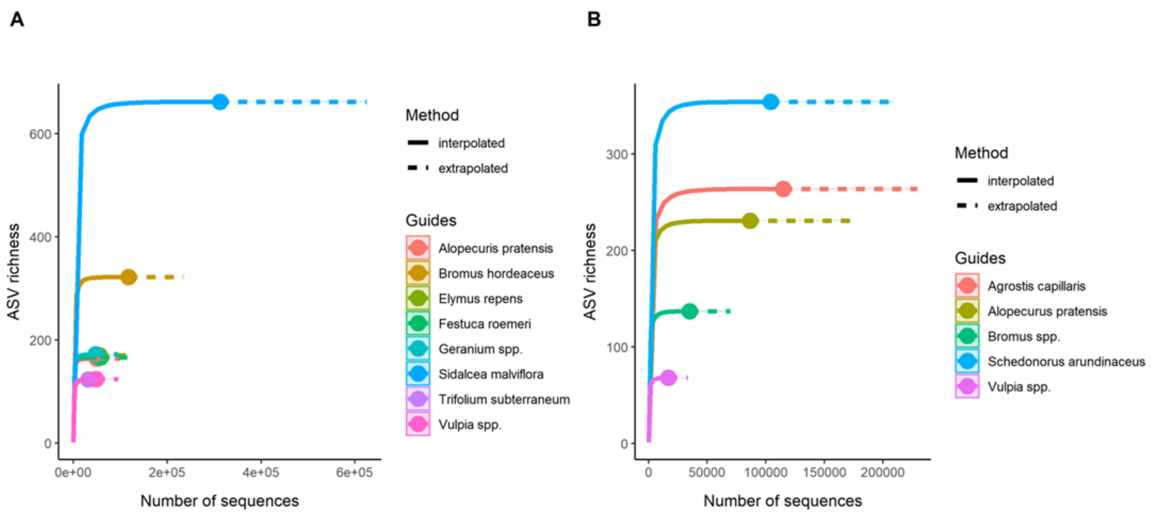


**Figure S1.** ASV accumulation curves by host plant species for high (A) and low (B) diversity plots across all sites. Curves were generated based on unnormalized ASV counts detected in roots of each host plant species (excluding plant species with <6 samples across sites). Rarefaction curves with extrapolation (solid and dashed lines, respectively) calculated with 200 replicate bootstraps for high- and low-diversity plots separately.

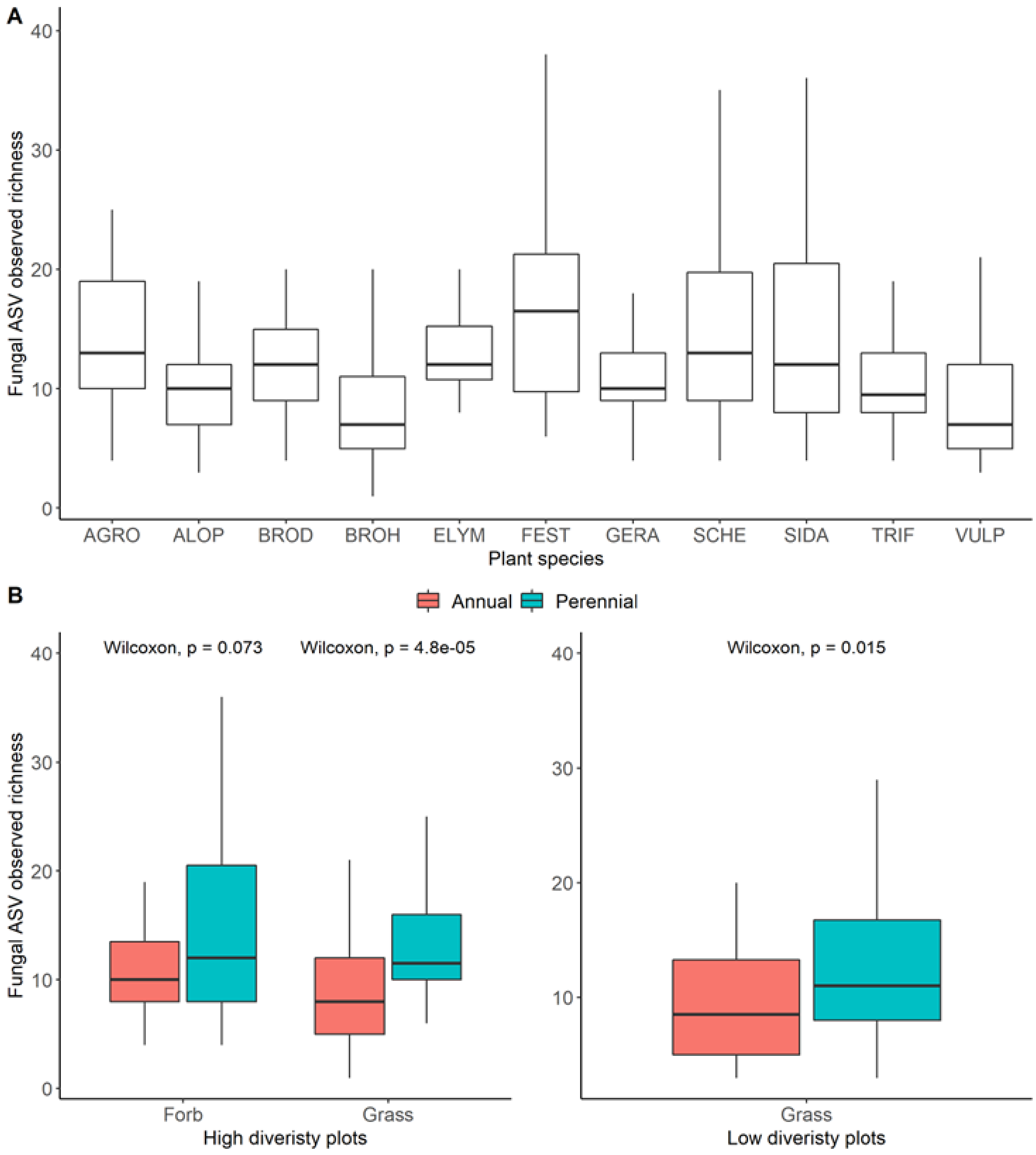


**Figure S2.** Boxplots depicting observed richness values for AM fungal ASVs detected across plant species (A) and plant functional groups in the high versus low diversity plots (B).

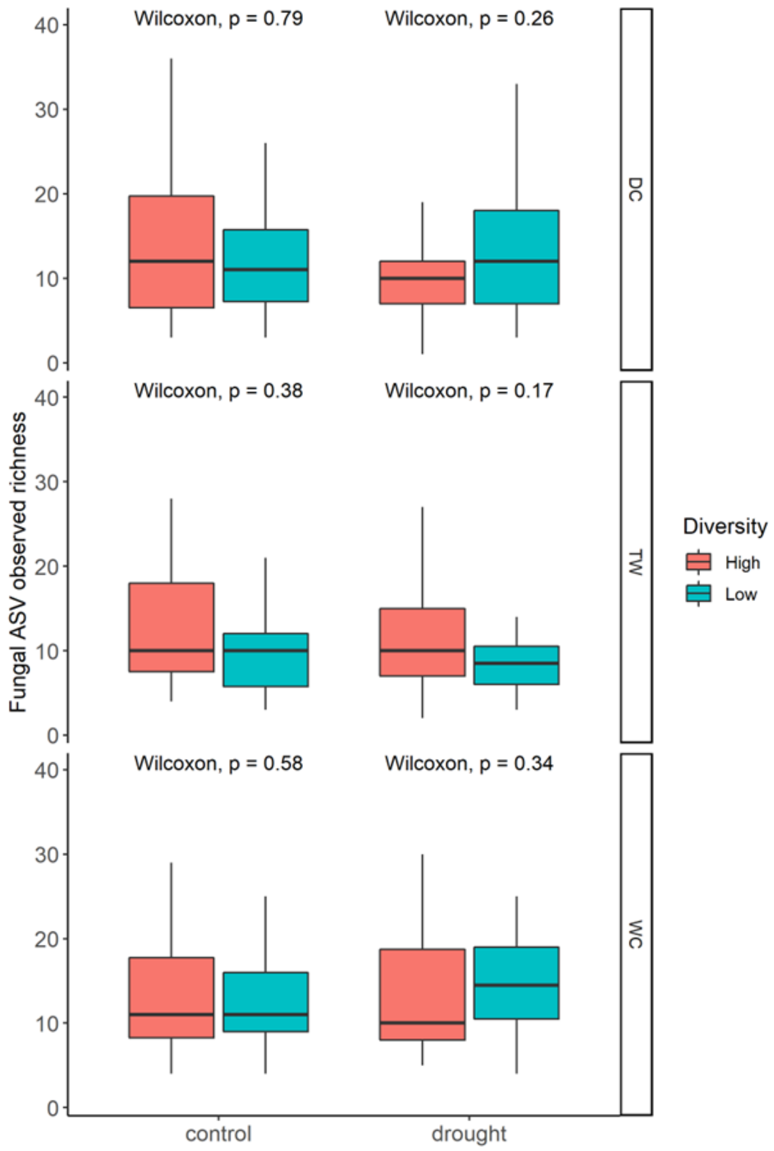


**Figure S3**. Boxplots depicting no effect of drought on the richness of AM fungal ASVs. No comparisons are significant.

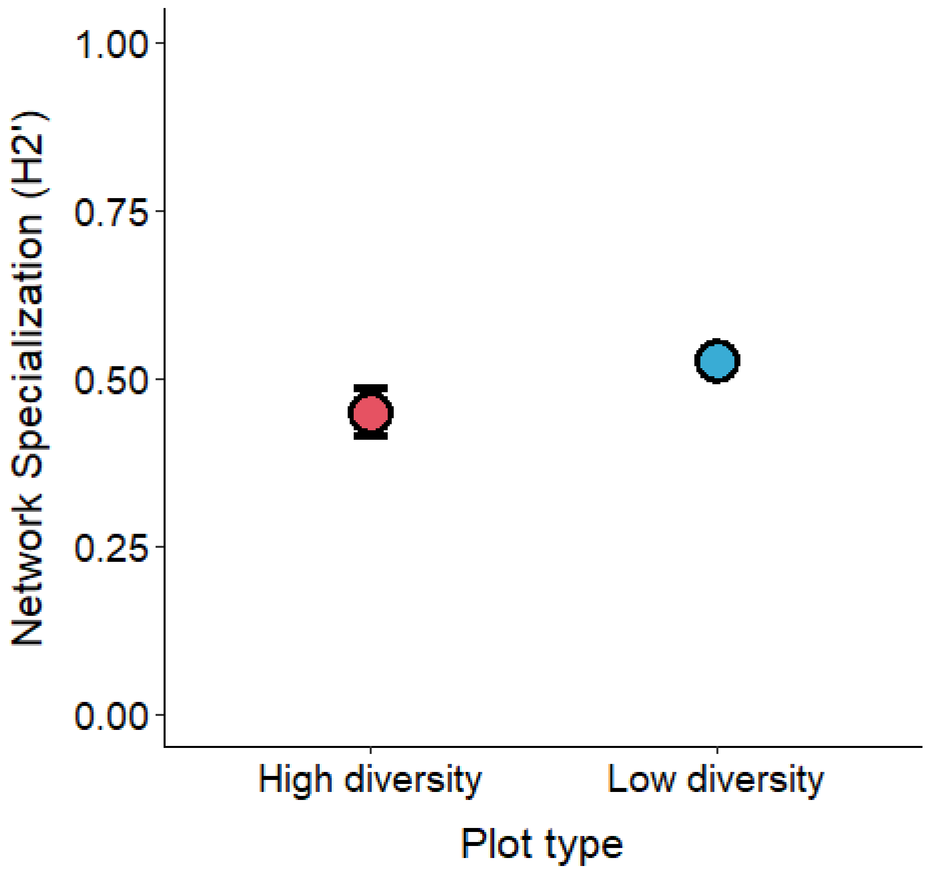


**Figure S4**. Overall network specialization (H2’) calculated for high and low diversity plots at each site revealed slightly more specialized plant-AM fungal networks for low diversity plots at the ASV level, but the relationship was not significant (Welch’s two sample t-test p=0.056, Bluthgen et al. 2006).

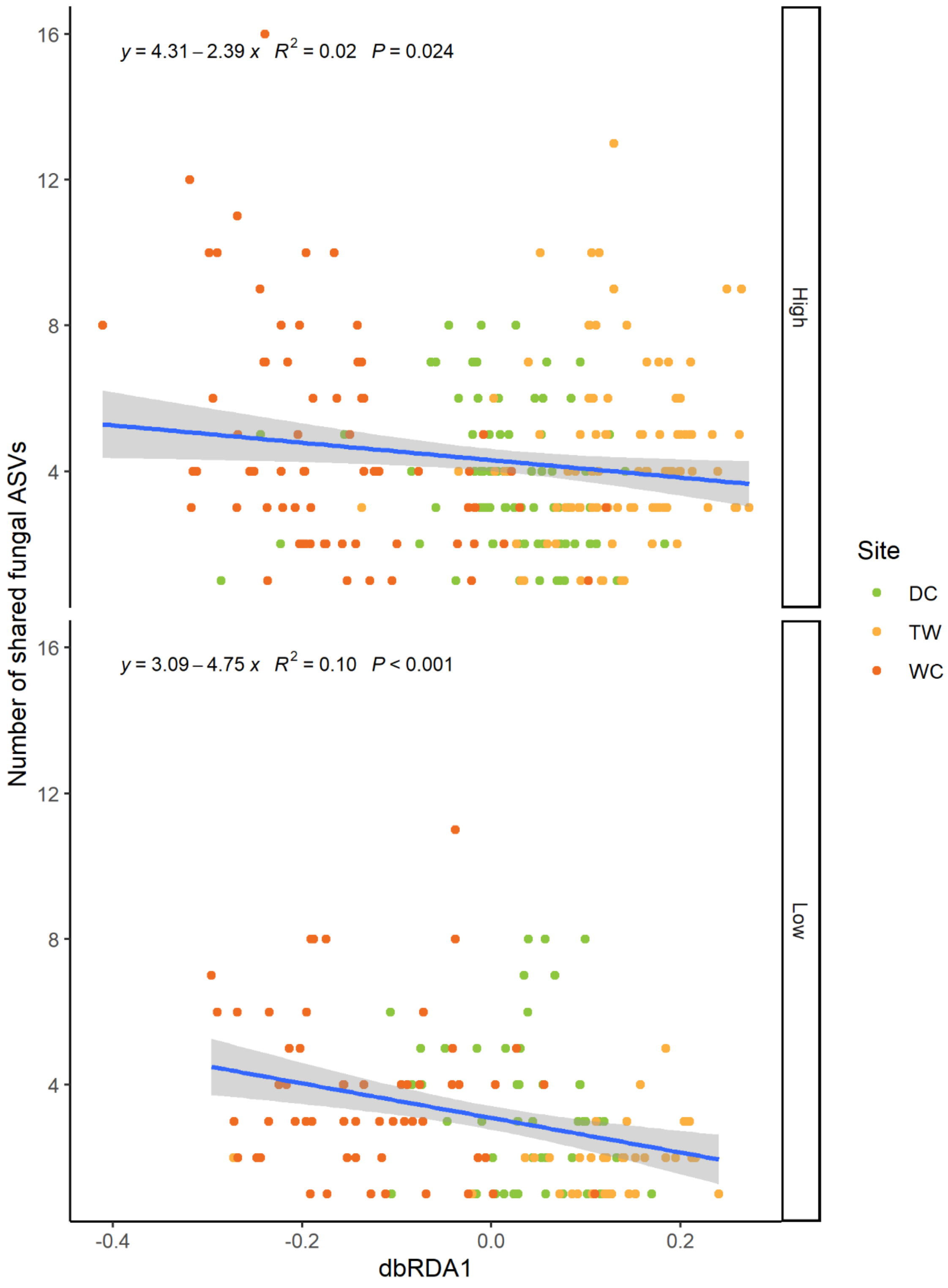


**Figure S5**. Significant declines in number of shared AM fungal ASVs between multiple plants within a plot and the first environmental axis from the dbRDA1 generated using soil nutrient and plant richness data.

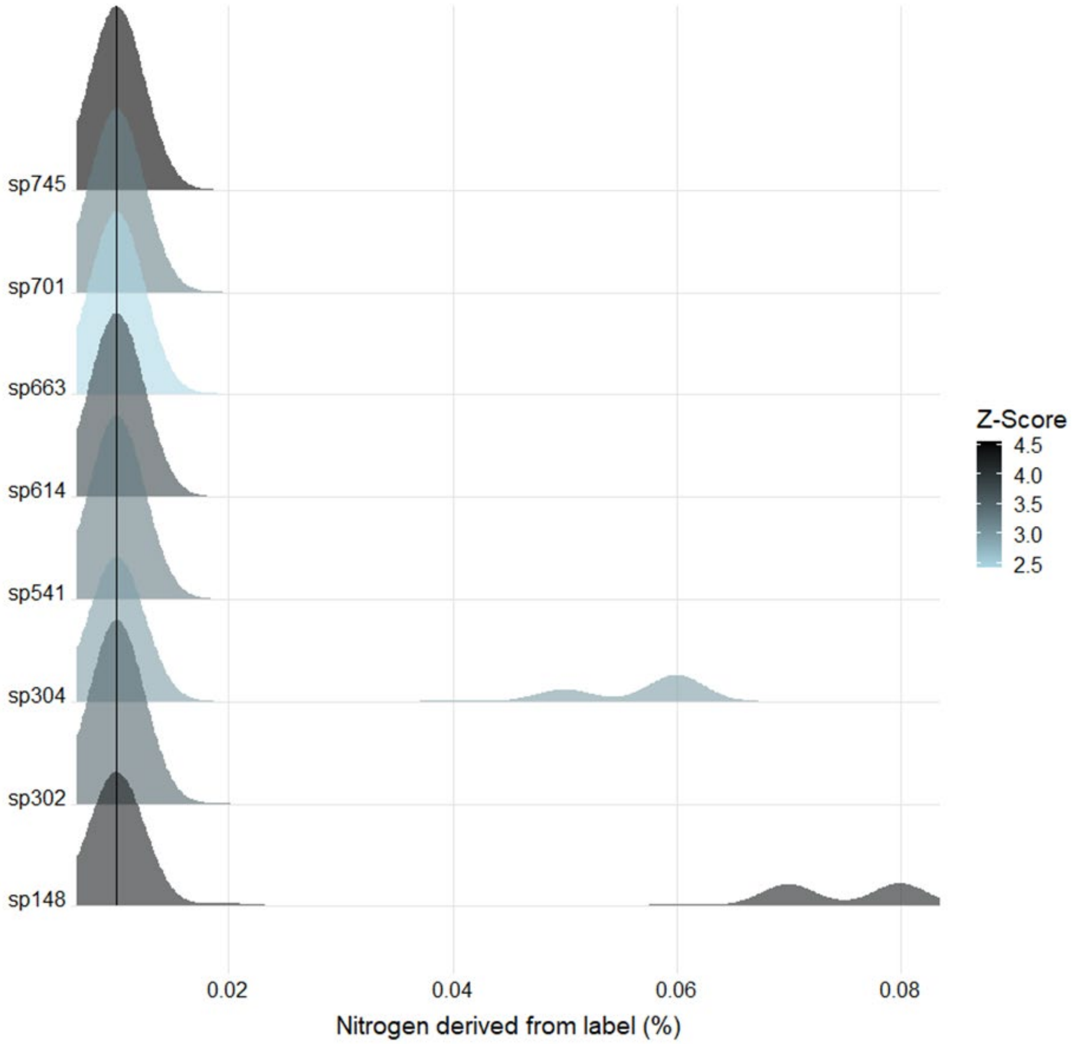


**Figure S6**. Results of threshold indicator taxa analysis (TITAN) for AM fungal ASVs that were indicators for declines in leaf 15N enrichment across all host plants.

**Table S1.** Module membership for AM fungal taxa in the high diversity plant-AM fungal interaction networks. C and z values are provided as the among-module connectivity (c) and within-module degree (z) columns.

| Family | Genus | Species | Module | Among-module connectivity | Within-module degree |
|---|---|---|---|---|---|
| unidentified | unidentified | unidentified | 2 | 0.652871844 | -0.136671164 |
| Paraglomeraceae | *Paraglomus* | VTX00001 | 1 | 0 | -0.526130062 |
| Archaeosporaceae | *Archaeospora* | VTX00004 | 4 | 0.674001256 | -1.391756482 |
| Archaeosporaceae | *Archaeospora* | VTX00005 | 4 | 0.710389499 | 0.605931665 |
| Archaeosporaceae | *Archaeospora* | VTX00008 | 1 | 0 | -0.526130062 |
| Archaeosporaceae | *Archaeospora* | VTX00009 | 1 | 0.734234117 | 0.054427248 |
| Acaulosporaceae | *Acaulospora* | VTX00021 | 1 | 0.71933866 | 2.231517158 |
| Acaulosporaceae | *Acaulospora* | VTX00023 | 4 | 0.708362525 | -1.140550188 |
| Acaulosporaceae | *Acaulospora* | VTX00026 | 1 | 0 | -0.526130062 |
| Acaulosporaceae | *Acaulospora* | VTX00027 | 1 | 0 | -0.526130062 |
| Acaulosporaceae | *Acaulospora* | VTX00028 | 1 | 0.624277719 | -0.235851407 |
| Acaulosporaceae | *Acaulospora* | VTX00030 | 1 | 0.601657861 | 2.812074468 |
| Gigasporaceae | *Gigaspora* | VTX00039 | 1 | 0 | -0.526130062 |
| Diversisporaceae | *Diversispora* | VTX00040 | 1 | 0 | -0.526130062 |
| Acaulosporaceae | *Acaulospora* | VTX00047 | 1 | 0 | -0.526130062 |
| Gigasporaceae | *Scutellospora* | VTX00049 | 1 | 0.655632472 | -0.09071208 |
| Gigasporaceae | *Scutellospora* | VTX00052 | 3 | 0.336293294 | -0.041603399 |
| Glomeraceae | *Glomus* | VTX00053 | 1 | 0.469556418 | -0.235851407 |
| Diversisporaceae | *Diversispora* | VTX00054 | 1 | 0 | -0.526130062 |
| Claroideoglomeraceae | *Claroideoglomus* | VTX00056 | 2 | 0.69350312 | 2.435463161 |
| Claroideoglomeraceae | *Claroideoglomus* | VTX00057 | 1 | 0.365420382 | 0.054427248 |
| Diversisporaceae | *Diversispora* | VTX00060 | 2 | 0.40937163 | -0.707124717 |
| Diversisporaceae | *Glomus* | VTX00061 | 1 | 0 | -0.235851407 |
| Glomeraceae | *Glomus* | VTX00064 | 2 | 0 | -0.640911358 |
| Glomeraceae | *Glomus* | VTX00067 | 1 | 0.698498721 | 2.66693514 |
| Glomeraceae | *Glomus* | VTX00070 | 1 | 0 | -0.526130062 |
| Glomeraceae | *Glomus* | VTX00072 | 2 | 0.595346684 | -0.663831367 |
| Glomeraceae | *Glomus* | VTX00074 | 4 | 0.577485165 | -0.15765572 |

| | | | | | |
|---|---|---|---|---|---|
| Glomeraceae | *Glomus* | VTX00080 | 1 | 0 | -0.526130062 |
| Glomeraceae | *Glomus* | VTX00086 | 2 | 0 | -0.707124717 |
| Glomeraceae | *Glomus* | VTX00093 | 1 | 0 | -0.526130062 |
| Glomeraceae | *Glomus* | VTX00096 | 1 | 0 | -0.526130062 |
| Glomeraceae | *Glomus* | VTX00101 | 1 | 0 | -0.526130062 |
| Glomeraceae | *Glomus* | VTX00108 | 2 | 0.708430649 | 2.522049861 |
| Glomeraceae | *Glomus* | VTX00113 | 4 | 0.556780934 | 0.440454503 |
| Glomeraceae | *Glomus* | VTX00114 | 4 | 0.198186757 | -1.391756482 |
| Glomeraceae | *Glomus* | VTX00115 | 4 | 0.562587334 | 0.066635613 |
| Glomeraceae | *Glomus* | VTX00125 | 3 | 0.47280272 | -0.762489889 |
| Glomeraceae | *Glomus* | VTX00126 | 1 | 0 | -0.526130062 |
| Glomeraceae | *Glomus* | VTX00135 | 2 | 0 | -0.707124717 |
| Glomeraceae | *Glomus* | VTX00137 | 2 | 0 | -0.707124717 |
| Glomeraceae | *Glomus* | VTX00142 | 3 | 0.502451908 | -0.873671387 |
| Glomeraceae | *Glomus* | VTX00143 | 4 | 0.706011089 | 0.472353715 |
| Glomeraceae | *Glomus* | VTX00155 | 2 | 0 | -0.707124717 |
| Glomeraceae | *Glomus* | VTX00159 | 3 | 0.328287594 | 0.723858161 |
| Glomeraceae | *Glomus* | VTX00160 | 3 | 0.533551558 | 0.955187407 |
| Glomeraceae | *Glomus* | VTX00166 | 3 | 0.295153396 | 0.205608918 |
| Glomeraceae | *Glomus* | VTX00180 | 1 | 0 | -0.526130062 |
| Glomeraceae | *Glomus* | VTX00183 | 1 | 0 | -0.526130062 |
| Glomeraceae | *Glomus* | VTX00191 | 1 | 0 | -0.526130062 |
| Claroideoglomeraceae | *Claroideoglomus* | VTX00193 | 2 | 0.675421507 | 1.353129411 |
| Glomeraceae | *Glomus* | VTX00197 | 2 | 0.570360047 | -0.289471223 |
| Glomeraceae | *Glomus* | VTX00199 | 1 | 0.672949237 | 2.376656486 |
| Glomeraceae | *Glomus* | VTX00204 | 1 | 0 | -0.526130062 |
| Glomeraceae | *Glomus* | VTX00209 | 1 | 0 | -0.526130062 |
| Glomeraceae | *Glomus* | VTX00212 | 2 | 0.708617471 | -0.447364617 |
| Glomeraceae | *Glomus* | VTX00214 | 2 | 0.40937163 | -0.707124717 |
| Glomeraceae | *Glomus* | VTX00216 | 4 | 0.48466135 | 2.107188326 |
| Glomeraceae | *Glomus* | VTX00219 | 3 | 0.660753413 | 2.11004039 |
| Glomeraceae | *Glomus* | VTX00222 | 3 | 0.329867931 | 0.659301162 |
| Glomeraceae | *Glomus* | VTX00223 | 3 | 0.3548493 | -0.809114388 |
| Claroideoglomeraceae | *Claroideoglomus* | VTX00225 | 1 | 0.664613732 | 1.796099176 |

| | | | | | |
|---|---|---|---|---|---|
| Acaulosporaceae | *Acaulospora* | VTX00228 | 3 | 0.209554787 | -0.873671387 |
| Acaulosporaceae | *Acaulospora* | VTX00230 | 3 | 0.226745091 | -0.237323825 |
| Acaulosporaceae | *Acaulospora* | VTX00231 | 1 | 0.70203254 | 0.48984523 |
| Claroideoglomeraceae | *Claroideoglomus* | VTX00237 | 2 | 0.678553507 | 0.365022363 |
| Archaeosporaceae | *Archaeospora* | VTX00245 | 4 | 0.723937932 | -0.707917126 |
| Glomeraceae | *Glomus* | VTX00256 | 1 | 0 | -0.526130062 |
| Claroideoglomeraceae | *Claroideoglomus* | VTX00276 | 3 | 0.663991082 | -0.753523639 |
| Claroideoglomeraceae | *Claroideoglomus* | VTX00278 | 2 | 0.542422969 | -0.024617787 |
| Claroideoglomeraceae | *Claroideoglomus* | VTX00279 | 3 | 0.38369548 | -0.873671387 |
| Paraglomeraceae | *Paraglomus* | VTX00281 | 4 | 0.720865849 | 0.798323787 |
| Glomeraceae | *Glomus* | VTX00295 | 1 | 0 | -0.526130062 |
| Glomeraceae | *Glomus* | VTX00296 | 1 | 0.484662773 | -0.235851407 |
| Diversisporaceae | *Diversispora* | VTX00306 | 1 | 0.525038004 | 1.650959849 |
| Glomeraceae | *Glomus* | VTX00309 | 4 | 0.479857889 | 0.589982059 |
| Glomeraceae | *Glomus* | VTX00315 | 3 | 0.434473087 | 1.573858648 |
| Archaeosporaceae | *Archaeospora* | VTX00338 | 2 | 0.464733254 | -0.37860459 |
| Claroideoglomeraceae | *Claroideoglomus* | VTX00340 | 1 | 0.244777575 | 0.054427248 |
| Diversisporaceae | *Diversispora* | VTX00380 | 2 | 0.483181707 | -0.312391232 |
| Glomeraceae | *Glomus* | VTX00399 | 1 | 0 | -0.526130062 |
| Claroideoglomeraceae | *Claroideoglomus* | VTX00402 | 2 | 0.670443149 | 0.871809225 |
| Glomeraceae | *Glomus* | VTX00410 | 1 | 0 | -0.526130062 |
| Glomeraceae | *Glomus* | VTX00423 | 4 | 0.667746638 | -0.29123367 |
| Glomeraceae | *Glomus* | VTX00426 | 1 | 0 | -0.526130062 |
| Paraglomeraceae | *Paraglomus* | VTX00444 | 2 | 0.61279728 | 0.296262336 |
| Glomeraceae | *Glomus* | VTX00445 | 2 | 0 | -0.707124717 |
| Paraglomeraceae | *Paraglomus* | VTX00446 | 1 | 0 | -0.526130062 |
| Archaeosporaceae | *Archaeospora* | VTX00456 | 3 | 0.532298644 | -1.002785385 |

**Table S2.** Module membership for AM fungal taxa in the low diversity plant-AM fungal interaction networks. C and z values are provided as the among-module connectivity (c) and within-module degree (z) columns.

| Family | Genus | Species | Module | Among-module connectivity | Within-module degree |
|---|---|---|---|---|---|
| unidentified | unidentified | unidentified | 3 | 0.389423452 | -0.077389947 |
| Archaeosporaceae | *Archaeospora* | VTX00004 | 2 | 0.413911778 | -0.774747707 |
| Archaeosporaceae | *Archaeospora* | VTX00005 | 2 | 0.623868615 | 0.258249236 |
| Archaeosporaceae | *Archaeospora* | VTX00009 | 3 | 0.327538423 | -0.700245592 |
| Acaulosporaceae | *Acaulospora* | VTX00021 | 3 | 0.478847723 | -0.01974778 |
| Acaulosporaceae | *Acaulospora* | VTX00023 | 2 | 0 | -0.860830785 |
| Acaulosporaceae | *Acaulospora* | VTX00028 | 2 | 0 | -0.860830785 |
| Acaulosporaceae | *Acaulospora* | VTX00030 | 1 | 0.554111376 | -0.385016805 |
| Gigasporaceae | *Scutellospora* | VTX00049 | 3 | 0.403705812 | -0.815529927 |
| Gigasporaceae | *Scutellospora* | VTX00052 | 1 | 0.258397448 | -0.158711366 |
| Glomeraceae | *Glomus* | VTX00053 | 3 | 0 | -0.077389947 |
| Claroideoglomeraceae | *Claroideoglomus* | VTX00056 | 1 | 0.560221621 | 2.738331695 |
| Claroideoglomeraceae | *Claroideoglomus* | VTX00057 | 1 | 0.480353059 | -0.68985361 |
| Diversisporaceae | *Diversispora* | VTX00060 | 3 | 0 | -0.873172095 |
| Glomeraceae | *Glomus* | VTX00064 | 3 | 0 | -0.653811624 |
| Glomeraceae | *Glomus* | VTX00067 | 3 | 0.306756952 | 0.037894388 |
| Glomeraceae | *Glomus* | VTX00072 | 1 | 0.368115343 | -0.070198485 |
| Glomeraceae | *Glomus* | VTX00074 | 2 | 0 | 1.463412335 |
| Glomeraceae | *Glomus* | VTX00079 | 2 | 0 | -0.860830785 |
| Glomeraceae | *Glomus* | VTX00086 | 1 | 0 | -0.916159049 |
| Glomeraceae | *Glomus* | VTX00089 | 2 | 0 | -0.860830785 |
| Glomeraceae | *Glomus* | VTX00108 | 2 | 0.617557776 | 1.807744649 |
| Glomeraceae | *Glomus* | VTX00113 | 2 | 0.652854882 | 0.430415393 |
| Glomeraceae | *Glomus* | VTX00115 | 2 | 0.665607992 | 1.31E-16 |
| Glomeraceae | *Glomus* | VTX00125 | 1 | 0.54763503 | -0.689009187 |
| Glomeraceae | *Glomus* | VTX00135 | 3 | 0 | -0.01974778 |
| Glomeraceae | *Glomus* | VTX00137 | 3 | 0 | -0.596169456 |
| Glomeraceae | *Glomus* | VTX00142 | 1 | 0 | -0.866056601 |
| Glomeraceae | *Glomus* | VTX00143 | 3 | 0.570322171 | 2.402824432 |
| Glomeraceae | *Glomus* | VTX00151 | 3 | 0 | -0.98845643 |
| Glomeraceae | *Glomus* | VTX00159 | 1 | 0.071237245 | 0.182847025 |
| Glomeraceae | *Glomus* | VTX00160 | 1 | 0.414982508 | 1.408798066 |
| Glomeraceae | *Glomus* | VTX00166 | 1 | 0.118803903 | 0.181158178 |
| Glomeraceae | *Glomus* | VTX00191 | 2 | 0 | -0.774747707 |
| Claroideoglomeraceae | *Claroideoglomus* | VTX00193 | 1 | 0.543114488 | 0.992086002 |

| | | | | | |
|---|---|---|---|---|---|
| Glomeraceae | *Glomus* | VTX00197 | 3 | 0.472236274 | -0.527319089 |
| Glomeraceae | *Glomus* | VTX00199 | 3 | 0.560880307 | -0.192674283 |
| Glomeraceae | *Glomus* | VTX00212 | 3 | 0.612848292 | 1.352456044 |
| Glomeraceae | *Glomus* | VTX00214 | 1 | 0.326338185 | -0.891107825 |
| Glomeraceae | *Glomus* | VTX00216 | 2 | 0.16330027 | 0.946913864 |
| Glomeraceae | *Glomus* | VTX00219 | 2 | 0.590034132 | 2.324243121 |
| Glomeraceae | *Glomus* | VTX00222 | 1 | 0.339440793 | 0.409996888 |
| Glomeraceae | *Glomus* | VTX00223 | 2 | 0 | -0.860830785 |
| Claroideoglomeraceae | *Claroideoglomus* | VTX00225 | 2 | 0.201446281 | -0.516498471 |
| Acaulosporaceae | *Acaulospora* | VTX00228 | 1 | 0.319408382 | -0.814265307 |
| Acaulosporaceae | *Acaulospora* | VTX00230 | 1 | 0.313641547 | -0.841005377 |
| Acaulosporaceae | *Acaulospora* | VTX00231 | 1 | 0.098896689 | 0.257156274 |
| Claroideoglomeraceae | *Claroideoglomus* | VTX00237 | 3 | 0.498908079 | 1.491757949 |
| Archaeosporaceae | *Archaeospora* | VTX00245 | 3 | 0.531696771 | 0.095536556 |
| Claroideoglomeraceae | *Claroideoglomus* | VTX00276 | 3 | 0.430584065 | 0.314897027 |
| Claroideoglomeraceae | *Claroideoglomus* | VTX00278 | 3 | 0.26807423 | 0.833676536 |
| Claroideoglomeraceae | *Claroideoglomus* | VTX00279 | 1 | 0 | -0.814265307 |
| Paraglomeraceae | *Paraglomus* | VTX00281 | 2 | 0.493931562 | 0.344332314 |
| Glomeraceae | *Glomus* | VTX00292 | 2 | 0 | -0.860830785 |
| Glomeraceae | *Glomus* | VTX00296 | 3 | 0 | -0.98845643 |
| Diversisporaceae | *Diversispora* | VTX00306 | 3 | 0.408906352 | 0.7183922 |
| Glomeraceae | *Glomus* | VTX00309 | 2 | 0 | 1.31E-16 |
| Glomeraceae | *Glomus* | VTX00310 | 3 | 0 | -0.98845643 |
| Glomeraceae | *Glomus* | VTX00315 | 1 | 0.368015671 | 1.737841671 |
| Paraglomeraceae | *Paraglomus* | VTX00335 | 3 | 0 | -0.98845643 |
| Archaeosporaceae | *Archaeospora* | VTX00338 | 3 | 0.475126429 | -0.135032115 |
| Claroideoglomeraceae | *Claroideoglomus* | VTX00340 | 1 | 0.420003393 | -0.81510973 |
| Diversisporaceae | *Diversispora* | VTX00380 | 3 | 0 | -0.307958618 |
| Claroideoglomeraceae | *Claroideoglomus* | VTX00402 | 1 | 0.612270242 | 0.042542849 |
| Glomeraceae | *Glomus* | VTX00423 | 2 | 0.514675627 | 0.516498471 |
| Paraglomeraceae | *Paraglomus* | VTX00444 | 3 | 0.502483716 | 2.633393103 |
| Glomeraceae | *Glomus* | VTX00445 | 3 | 0 | -0.930814263 |
| Archaeosporaceae | *Archaeospora* | VTX00456 | 2 | 0.476074219 | -0.860830785 |
